# Aquaporin-4 expression levels and mis-localization are inversely linked to peritumoral edema in gliomas of varying aggressiveness

**DOI:** 10.1101/2024.09.15.613106

**Authors:** Onofrio Valente, Pasqua Abbrescia, Gianluca Signorile, Raffaella Messina, Luigi de Gennaro, Maria Teresa Bozzi, Giuseppe Ingravallo, Antonio D’amati, Domenico Sergio Zimatore, Claudia Palazzo, Grazia Paola Nicchia, Antonio Frigeri, Francesco Signorelli

## Abstract

Aquaporin-4 (AQP4) and its extended isoform, AQP4ex, are crucial for regulating brain water homeostasis. Mis-localization of these isoforms is implicated in various brain tumors, including glioblastoma multiforme (GBM).

This study explores AQP4 isoform expression and localization in Pilocytic Astrocytomas (PA), a circumscribed astrocytic low-grade glioma, compared to GBM, an adult-type diffuse high grade glioma.

We found significant upregulation of AQP4 and AQP4ex in PA, with notable mis-localization deviating from the typical perivascular localization seen in healthy tissue. This mis-localization mirrors the phenotype observed in AQP4ex knockout models, where impaired AQP4 localization is linked to disrupted water homeostasis and reduced waste clearance, despite overall increased AQP4 levels. Interestingly, PA shows minimal peritumoral edema and a relatively intact blood-brain barrier (BBB), with elevated phosphorylated AQP4ex (pAQP4ex) suggesting a role in stabilizing AQP4 function. In contrast, GBM exhibits reduced AQP4/AQP4ex expression, significant peritumoral edema, and BBB disruption.

GFAP isoforms, GFAPκ and GFAPδ, are upregulated in PA, associated with Rosenthal fibers, indicating a stabilizing astrocytic response. GBM, however, shows generalized GFAP increase, reflecting aggressive gliosis and disrupted water homeostasis.

In conclusion, both AQP4 expression levels and mis-localization are important factors influencing peritumoral edema and tumor aggressiveness in gliomas. This study positions AQP4 as a potential biomarker for glioma progression, offering insights into astrocytic function and paving the way for targeted therapies.

## INTRODUCTION

Several previous reports [41–44] including our recent research [28, 29, 37], have demonstrated a significant involvement of aquaporin-4 (AQP4) in Glioblastoma multiforme (GBM). These studies have highlighted that the expression and distribution of AQP4 are altered in GBM, influencing peritumoral edema formation and tumor progression. The readthrough isoform, AQP4ex, plays a central role in both physiological AQP4 perivascular localization and mis-localization during GBM development. Indeed, we found that AQP4ex expression was almost absent in tumoral regions, while the canonical isoforms of AQP4 appeared primarily delocalized. On the other hand, in peritumoral samples AQP4 expression was altered in perivascular astrocyte processes, where AQP4ex was reduced and partially delocalized. Additionally, the severity of edema correlated with the downregulation of AQP4ex, suggesting a link between the reduction of expression of this isoform and the increased in peritumoral edema. In light of these results, it is of great interest to extend the analysis to other brain tumors, particularly pilocytic astrocytomas (PA). PA represent a significantly different class of brain tumors compared to GBM in terms of aggressiveness, progression potential, and prognosis [4, 25]. PA are grade 1 slow-growing circumscribed astrocytomas and generally have a favorable prognosis according to the 2021 World Health Organization (WHO) classification of central nervous system (CNS) tumors [16]. PA represent about 5-6% of all brain tumors and are the most common glial tumors in children, with a peak incidence between 5 and 15 years of age. Their typical localization in the cerebellum can cause symptoms such as ataxia, dysmetria, and balance disturbances [20]. Other locations, such as the brainstem and spinal cord, can cause a variety of neurological symptoms depending on the affected area [3]. Peritumoral edema is a significant complication in both GBM and PA as it can significantly impact the patient’s quality of life and the clinical management of the tumor. Edema can increase intracranial pressure and cause additional neurological symptoms, requiring careful and timely management. However, its management can differ greatly due to the distinct biological characteristics of the two tumors. Understanding whether AQP4 and AQP4ex influence tumor growth and peritumoral edema differently in PA compared to GBM, can reveal unique biological mechanisms that could potentially be exploited for new therapies.

The aim of the present study is to extend the knowledge acquired on the role of AQP4 and AQP4ex in GBM to PA. Specifically, we aim to evaluate the expression levels and localization of AQP4 and AQP4ex in PA, to determine their role in the formation of peritumoral edema in PA, and compare the results obtained with those previously observed in GBM to identify similarities and differences in the regulatory mechanisms of AQP4 between these two types of brain tumors. Furthermore, for the first time we evaluate phosphorylation level of AQP4ex (pAQP4ex) in PA and GBM samples and correlate these with AQP4 expression levels.

## MATERIAL AND METHODS

### TISSUE COLLECTION AND HISTOLOGICAL CHARACTERIZATION

#### Tissue Collection and Histological Characterization

**Samples** from PA and GBM were obtained during neurosurgical tumor resections. Seven GBM samples (Pts. #5, 6, 7, 8, 9, 16, 17) used for this study were previously described in our earlier research [37].

Tumor (T) samples corresponded to tissue clearly infiltrated by PA or GBM on microscopic vision under white light and confirmed by neuronavigation. Non-infiltrated samples (N) were defined as anatomopathologically-verified normal brain parenchyma surrounding the tumor, supplied together with T samples from the same patient and were used as controls for GBM T samples, as previously reported [37].Unlike GBM, for whom the current standard, if functionally feasible, is supratotal resection that may include non-infiltrated brain regions [2, 23] used as controls, for PA surgical resection is generally limited to tumor-infiltrated brain parenchyma. Thus, we used as controls of PA-infiltrated samples microscopically confirmed healthy tissue obtained from different patients during surgery for cerebellar pathologies.

### MAGNETIC RESONANCE IMAGING (MRI) AND EDEMA INDEX CALCULATION

All images were acquired using a 1.5 Tesla MRI scanner equipped with a standard coil. An expert neuroradiologist (DSZ) quantified the volume of tumor and relative peritumoral edema in the PA patients. The image datasets comprised: 1) T1-weighted, gadolinium enhanced (T1c) images with 3D acquisition and 1mm isotropic voxel sizes for all patients, and 2) FLAIR images with 3D acquisition and 1mm isotropic voxel sizes or T2-weighted images with 2D axial acquisition and 5mm thickness. T1c images were used for tumor segmentation, while FLAIR/T2 sequences were utilized to delineate edema and non-enhancing tumor components.

Tumor and edema volumes were calculated using 3D Slicer Software (Release 4.10.2). T1c images were co-registered to FLAIR/T2 images. An interactive segmentation algorithm was employed: solid tumor portions and cystic portions were delineated “slice by slice” on T1c images, followed by manual corrections. Edema and non-enhancing tumor components were manually segmented similarly. The solid tumor portions, both enhancing and non-enhancing, and the cystic portion were defined as the “overall tumor volume. The Edema Index (EI), calculated as tumor volume + edema volume/tumor volume, was used to estimate the extent of peritumoral edema in patients with PA [13, 19, 37]. Two EI categories were identified based on the evaluation of MRI images: EI = 1 (edema absent); EI >1 (edema present).

### ANTIBODIES

For the immunoblot and immunofluorescence experiments the following primary antibodies were used: custom anti p-AQP4ex antibody at a concentration of 0.2 µg/mL for immunoblot and at 0.4 µg/mL for immunofluorescence[22]. A rabbit anti-human AQP4ex [7], used at a concentration of 0.3 μg/mL and a rabbit anti-AQP4 antibody [37] was used at a concentration of 0.1 μg/mL for immunoblot and at 0.4 μg/mL for immunofluorescence. Therefore, Mouse anti-vascular endothelial growth factor (VEGF) was used at 2 μg/mL and custom mouse Anti-GFAP antibody (Sigma-Aldrich, Cat# IF03L) was used at concentration of 0.1 μg/mL for both immunoblot and immunofluorescence. The secondary antibody used for immunoblotting experiments were anti-rabbit IgG-HRP (Bio-Rad Cat# 172-101) and anti-mouse IgG-HRP (Bio-Rad Conjugate #1706516) and Alexa Fluor 488 anti-rabbit (Life Technologies, Cat# A-11034) and Alexa Fluor 488 anti-mouse (Life Technologies, Cat # A28175) for immunofluorescence, used at a concentration of 1 μg/mL.

### SAMPLE PREPARATION FOR SDS-PAGE AND IMMUNOBLOT

Tissues for experiments were prepared as previously reported [37].

PA, GBM and N regions were dissolved in seven volumes of BN buffer (1% Triton X-100, 12 mM NaCl, 500 mM 6-aminohexanoic acid, 20 mM Bis-Tris, pH 7.0; 2 mM EDTA; 10% glycerol) with a protease inhibitor cocktail (Roche Diagnostic, Monza, Italy). Tissue lysis was performed on ice for 1 hour, and the samples were then centrifuged at 17,000 × g for 30 minutes at 4 °C. Supernatants were collected, and total protein content was calculated using the BCA Protein Assay Kit (Pierce-Thermo Fisher Scientific, USA). For SDS-page experiments 25 μg of homogenates were dissolved in Laemmli sample buffer 2X (Bio-Rad, California, USA) added with 50 mM dithiothreitol (DTT) and proteins were separated on 13% polyacrylamide gel and transferred to polyvinylidene difluoride (PVDF) membranes (Millipore, Burlington, MA, USA) for immunoblot analysis. PVDF membranes were incubated as previously reported [37].

Reactive proteins were revealed using chemiluminescent detection system (Clarity Western ECL Substrate, Bio-Rad, California, USA) and visualized on a Chemidoc Touch imaging system (Bio-Rad, California, USA). Densitometry analysis was performed using Image Lab (Bio-Rad California, USA), and the relative expression of proteins was normalized with Rouge Ponceaux staining. The total amount of p-AQP4ex, AQP4ex and global AQP4 expression of N and T tissues was represented as the percentage change from N control tissues, set on 100%. Finally, the percentage of p-AQP4ex was also represented as ratio (pAQP4ex/AQP4ex) indicating the change in the phosphorylated isoform relative to the total extended isoforms.

### IMMUNOFLUORESCENCE

Immunofluorescence experiments were performed as previously described [37]. Ten μm fresh cryosections of each sample were briefly fixed in 4% PFA solution, and after blocking (PBS-Gelatin 0.1%, 15 minutes at room temperature (RT)) were incubated with primary antibodies (1h, RT). Before incubation with secondary antibody (1h, RT) cryosections were washed for 15 minutes with blocking solution and then mounted with Mowiol (Sigma-Aldrich) added with DAPI (4′,6-diamidino-2-phenylindole, Life Technologies, Thermo Fisher Scientific, Waltham, Massachusetts, USA). Finally, images were obtained under an SP8 confocal automated inverted Lightning microscope (Leica TCS) using 20×/0.55 HC PL FLUOTAR objectives or with a 63X HC PL Apo oil CS2 objective.

### EVALUATION OF SODIUM FLUORESCEIN (SF) CONCENTRATION

Sodium Fluorescein (SF) (Monico Spa, Italy) is well-documented for guiding the resection of high and lower-grade gliomas, including GBM [1, 2, 23] and PA. According to our protocol, SF was administered intravenously at a dose of 5 mg/kg at the induction of anesthesia[9]. SF concentration was measured in GT samples from PA and GBM patients and in N samples from GBM from GBM patients, consistent with our previous study [37]. Fluorescence was assessed using an automatic microplate reader (FLEX Station, Molecular Devices) and for that procedure, 40 μL of each sample homogenates, was placed in 96-well black-walled, clear-bottom microplates (Corning, NY, USA), and fluorescence was measured using SoftMax Pro software.

### STATISTICAL ANALYSIS

Immunoblot data and SF concentration are reported as a bar chart with the SEM. Statistical analysis was performed using GraphPad Prism 8 (GraphPad, San Diego, CA, USA). Data distribution was analyzed using the Shapiro-Wilk test: on parametric data the analysis of variance (One-Way ANOVA) followed by Tukey’s post-test was performed, while on non-parametric ones One-Way ANOVA on ranks (Kruskal-Wallis test) followed by Dunn’s multiple comparisons test was performed. For data set containing only two data set, student T-test (Welch’s t-test, for data that shows great differences in Standard Deviation (SD)) was performed for parametric data, and Mann-Whitney test was used for non-parametric data set. A p-value < 0.05 was considered statistically significant.

This study received approval from the local institutional review board (project. n.6898) and was conducted in accordance with the ethical principles for medical research involving human subjects outlined in the Declaration of Helsinki and its subsequent amendments.

## RESULTS

From a prospectively maintained institutional database including brain gliomas, 11 PA patients were included. Their demographic, radiological and histopathological data are listed in Table 1.

**Table 1.**
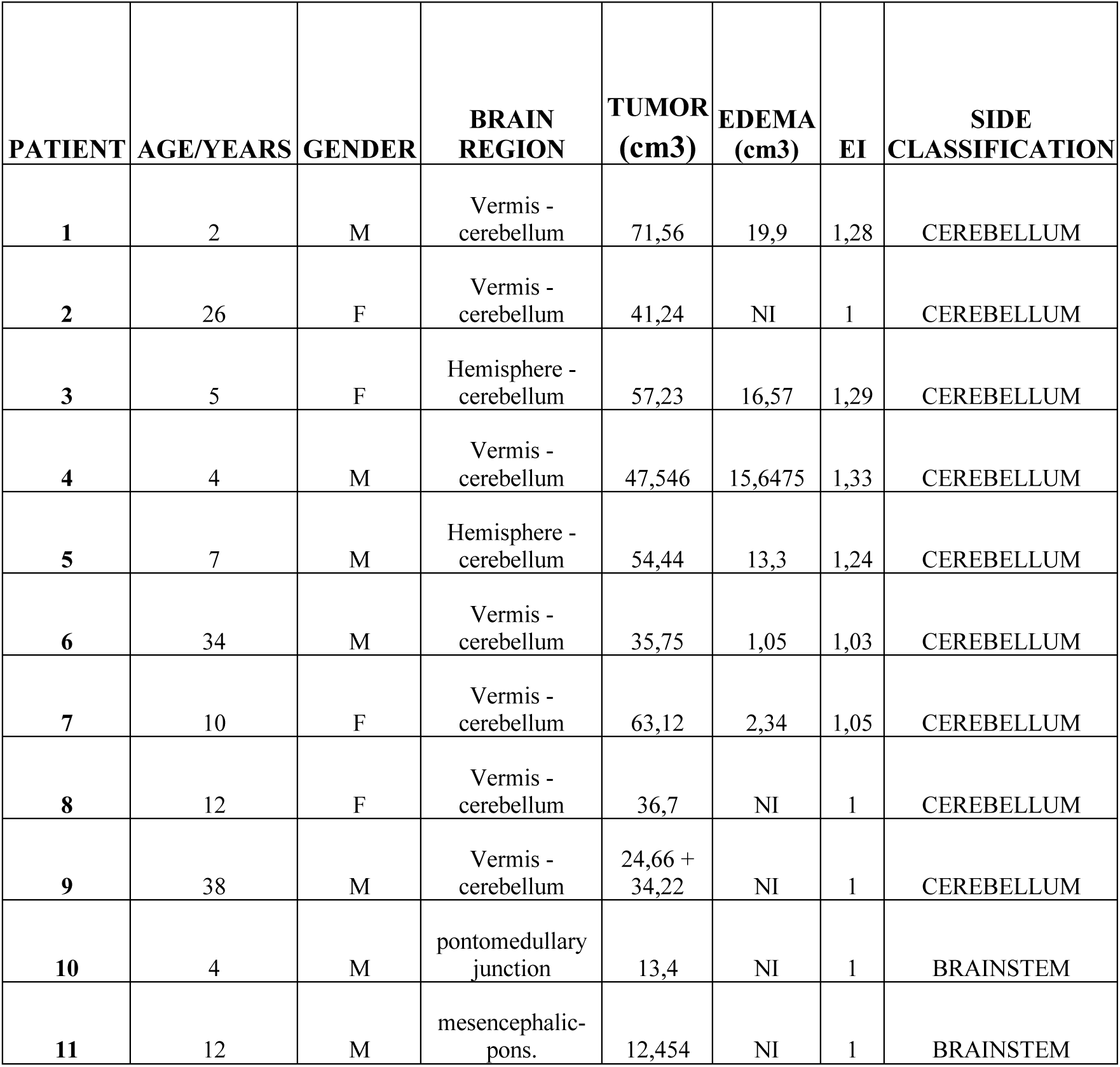
Features of the PA patient’s cohort.

| PATIENT | AGE/YEARS | GENDER | BRAIN REGION | TUMOR (cm3) | EDEMA (cm3) | EI | SIDE CLASSIFICATION |
| --- | --- | --- | --- | --- | --- | --- | --- |
| 1 | 2 | M | Vermis - cerebellum | 71,56 | 19,9 | 1,28 | CEREBELLUM |
| 2 | 26 | F | Vermis - cerebellum | 41,24 | NI | 1 | CEREBELLUM |
| 3 | 5 | F | Hemisphere - cerebellum | 57,23 | 16,57 | 1,29 | CEREBELLUM |
| 4 | 4 | M | Vermis - cerebellum | 47,546 | 15,6475 | 1,33 | CEREBELLUM |
| 5 | 7 | M | Hemisphere - cerebellum | 54,44 | 13,3 | 1,24 | CEREBELLUM |
| 6 | 34 | M | Vermis - cerebellum | 35,75 | 1,05 | 1,03 | CEREBELLUM |
| 7 | 10 | F | Vermis - cerebellum | 63,12 | 2,34 | 1,05 | CEREBELLUM |
| 8 | 12 | F | Vermis - cerebellum | 36,7 | NI | 1 | CEREBELLUM |
| 9 | 38 | M | Vermis - cerebellum | 24,66 + 34,22 | NI | 1 | CEREBELLUM |
| 10 | 4 | M | pontomedullary junction | 13,4 | NI | 1 | BRAINSTEM |
| 11 | 12 | M | mesencephalic-pons. | 12,454 | NI | 1 | BRAINSTEM |

### Pilocytic astrocytomas (PA) characterization: main histological features

The key histological characteristics of PA are showed in Figure 1, including glomeruloid microvascular proliferations (Fig. 1a), blood vessels with hyalinized walls and Rosenthal fibers in astocytes (Fig. 1b). Strong immunoperoxidase staining of GFAP (Fig. 1c) and and Oligodendrocyte Transcription Factor (OLIG2, Fig 1d)) confirmed the glial nature of PA [30, 34].

**Fig. 1.**
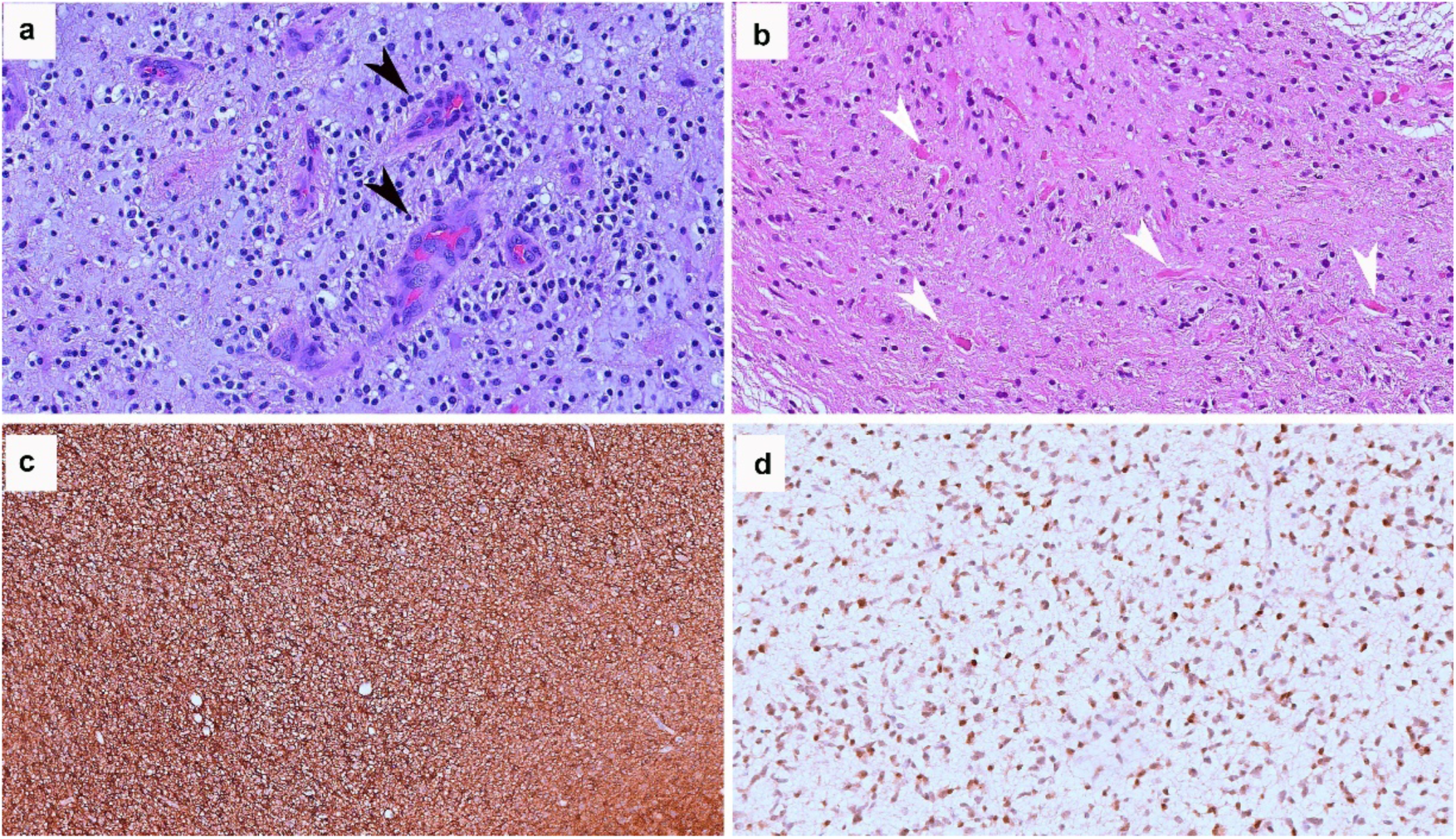
Hallmark histological features of pilocytic astrocyoma. (a-b) H&E-stained section of a pilocytic astrocytoma (Pt.6 in the table) (a) Glomeruloid microvascular proliferations (black arrows) (HE, 200x). (b) Numerous eosinophilic Rosenthal fibers in a compact fibrillary area (white arrows) (HE, 200x). (c-d) Immunohistochemical expression of glial markers in pilocytic astrocytoma. (c) Diffuse cytoplasmic positivity for GFAP (brown staining) in neoplastic elements (100x). (d) Nuclear expression of OLIG2 in tumor cells (nuclei stained in brown) (200x)

### AQP4 localization is affected in histologically characterized PA samples

We then investigated the expression levels and localization of AQP4 and AQP4ex in PA biopsies using immunofluorescence (Fig.2). The analysis involved also the phosphorylated form of AQP4ex (p-AQP4ex) using a specific antibody recognizing phosphorylated serine residue (Ser-335) previously identified [7] and recently demonstrated to be expressed in human brain [22]. N samples (CTRL) showed the perivascular staining for AQP4 (Fig. 2d), AQP4ex (Fig. 2g) and p-AQP4ex (Fig. 2j) due to astrocyte end-feet localization of AQP4. AQP4 staining was weak in PA samples and resulted not confined at perivascular level, but mainly appears distributed in the parenchyma (Fig. 2e). Interestingly, both AQP4ex (Fig. 2h) and p-AQP4ex (Fig. 2k) staining appeared also strongly reduced at the perivascular level and emerged sparsely diffused in the parenchyma.

In GBM T samples, AQP4 (Fig. 2f) as well as AQP4ex staining (Fig. 2i) were mis-localized and weak confirming what previously reported [37]. p-AQP4ex IF signal staining (Fig. 2l), was barely detectable and even less intense at the perivascular pole.

**Fig. 2.**
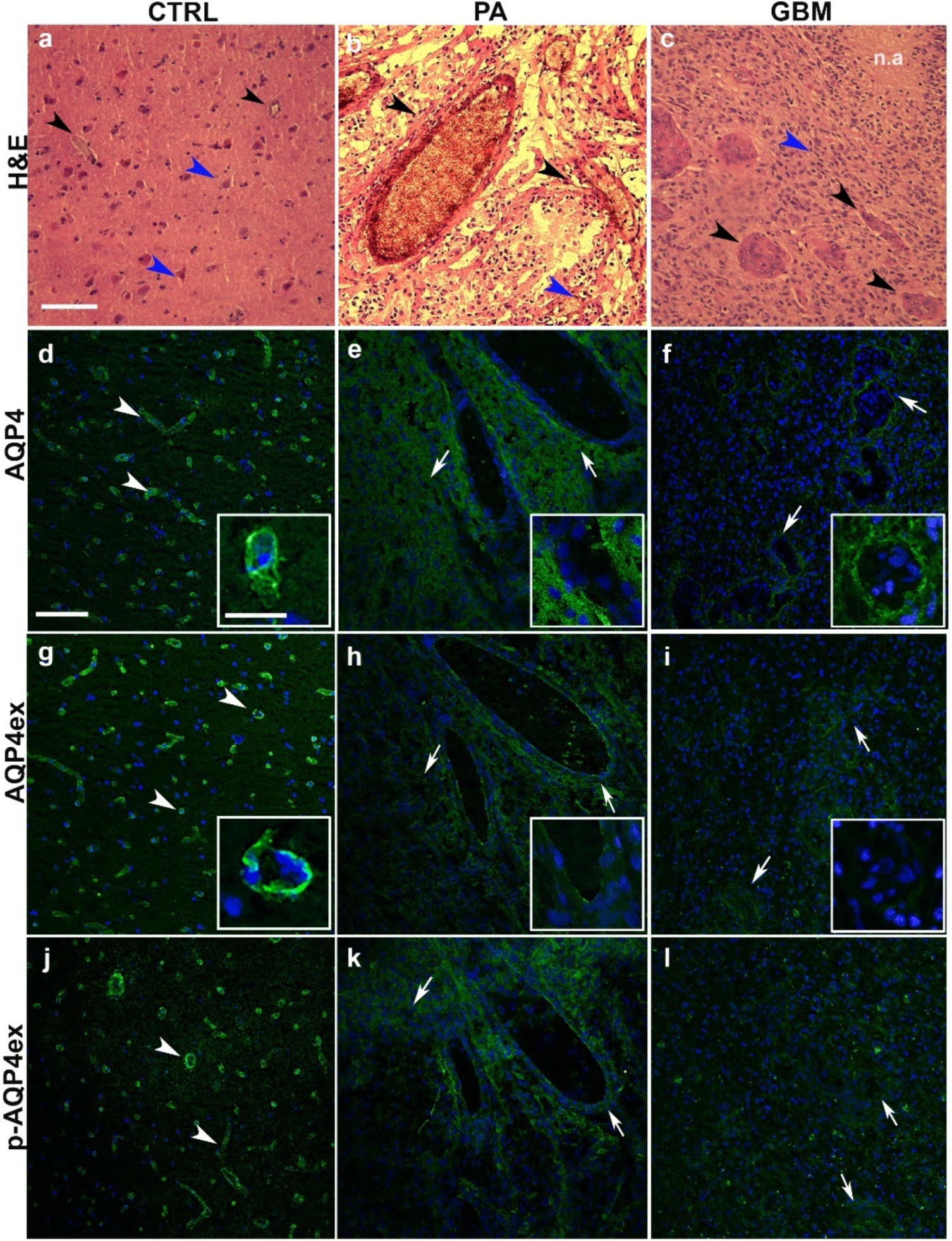
AQP4 isoform localization in PA and GBM. (a-c) Hematoxylin-eosin (H&E) staining of PA and GBM T (tumor) samples; (a) Typical section of N (CTRL, a), PA (b) and GBM (c); Cellularity in each tissue is indicated by blue arrowheads while blood vessels are indicated in each sample with white arrowheads. Necrotic areas (n.a) exclusive GBM’s feature is indicated in c. Scale bar 300 μm. (d-l) Immunofluorescence of AQP4 isoforms in N (CTRL), PA (Pt.1) and GBM-T (Pt.5) samples. AQP4 (d-f) and AQP4ex (g-i) and p-AQP4ex (j-l) staining in PA and GBM-T samples compared to N samples. Arrowheads indicates the blood vessels in CTRL tissue, while arrows show a weak perivascular signal in PA and GBM samples. Scale bar 300 μm. Inset scale bar 20 μm

### AQP4 isoforms protein expression is differently affected in PA and GBM

To quantitatively evaluate AQP4 isoforms levels, immunoblot experiments were performed in PA, normal cerebellar tissue, GBM T and N samples. Figure 3 shows immunoblot and densitometric analysis results for total AQP4, AQP4ex and p-AQP4ex expression levels in PA. Notably, the total amount of AQP4 (Fig 3b, right) of AQP4ex (Fig 3c, right), and the phosphorylated counterpart (Fig 3d, right) were significantly increased. More specifically, the increased p-AQP4ex/AQP4ex ratio confirmed that the increase of phosphorylation accounted for most of the observed increase of AQP4ex isoform content (Fig. 3e).

**Fig. 3.**
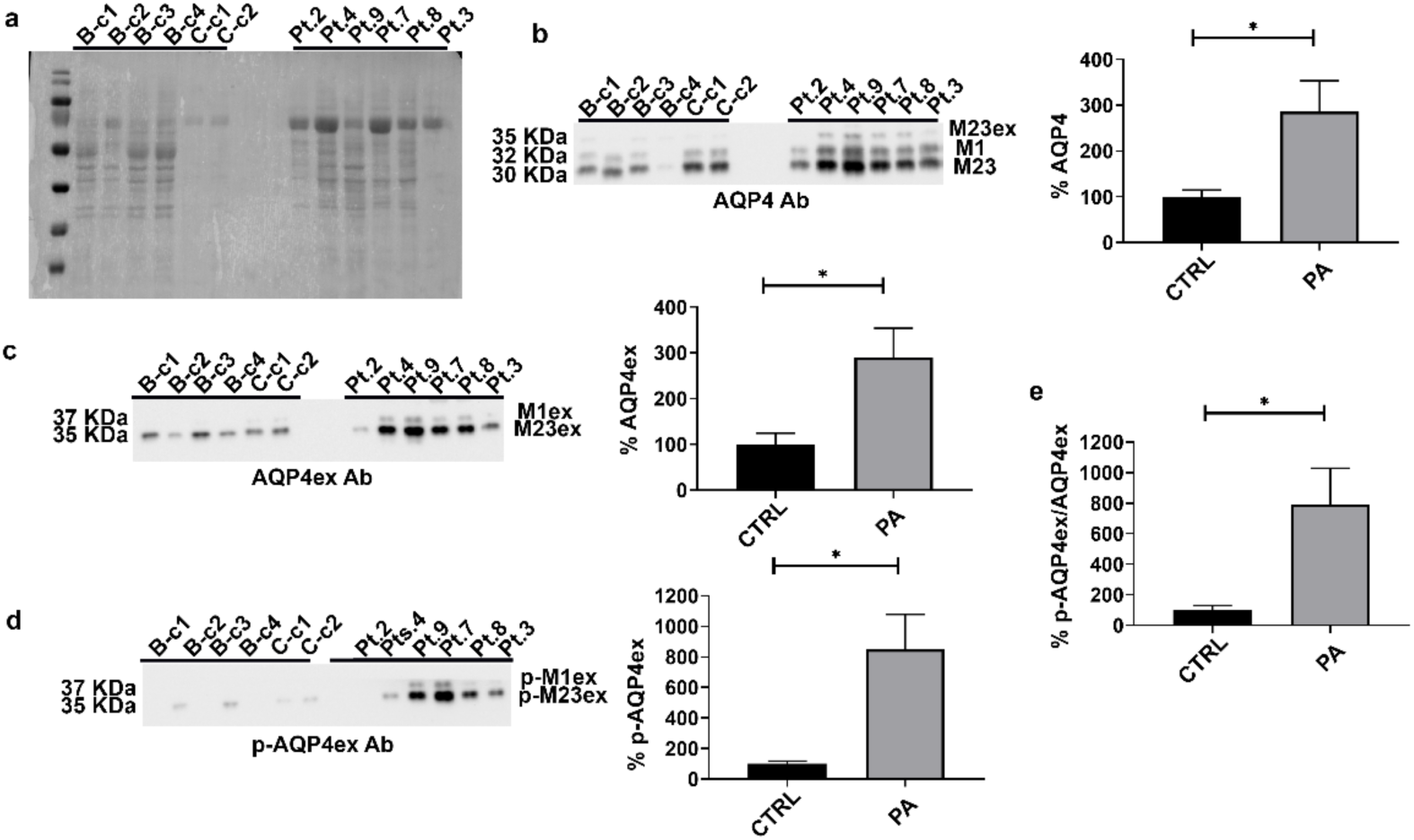
Quantitative analysis of AQP4 isoforms in PA tumors. (a) Rouge Ponceaux staining of total proteins after transfer onto a PVDF membrane. (b-c-d) Immunodetection (left) and densitometric analysis (right) of AQP4 isoforms using antibodies for global AQP4 (b), AQP4ex (c), and p-AQP4ex (d), respectively. (e) Ratio between p-AQP4ex and AQP4ex shows a strong increase of phosphorylation levels in PA compared to CTRL. (Asterisk indicated the student’s t-test significant differences for the comparison with basal level, *p < 0.05;) Brain CTRL (B-c1,2,3,4, n=4), Cerebellum CTRL (C-c1,2, n=2), PA (n=6)

In GBM T samples we confirmed the reduction of total AQP4 protein and of the AQP4ex isoform in the analyzed GBM sample, as previously reported [37]. Interestingly, differently to what found in PA, pAQP4ex was significantly reduced in GBM (Fig. 4d). In addition, the p-AQP4ex/AQP4ex ratio was found increased, (Fig. 4e), underlining that in GBM T samples, differently to PA, a decoupling occurs between the amount of AQP4ex produced and its levels of phosphorylation (Fig. 4e).

**Fig. 4.**
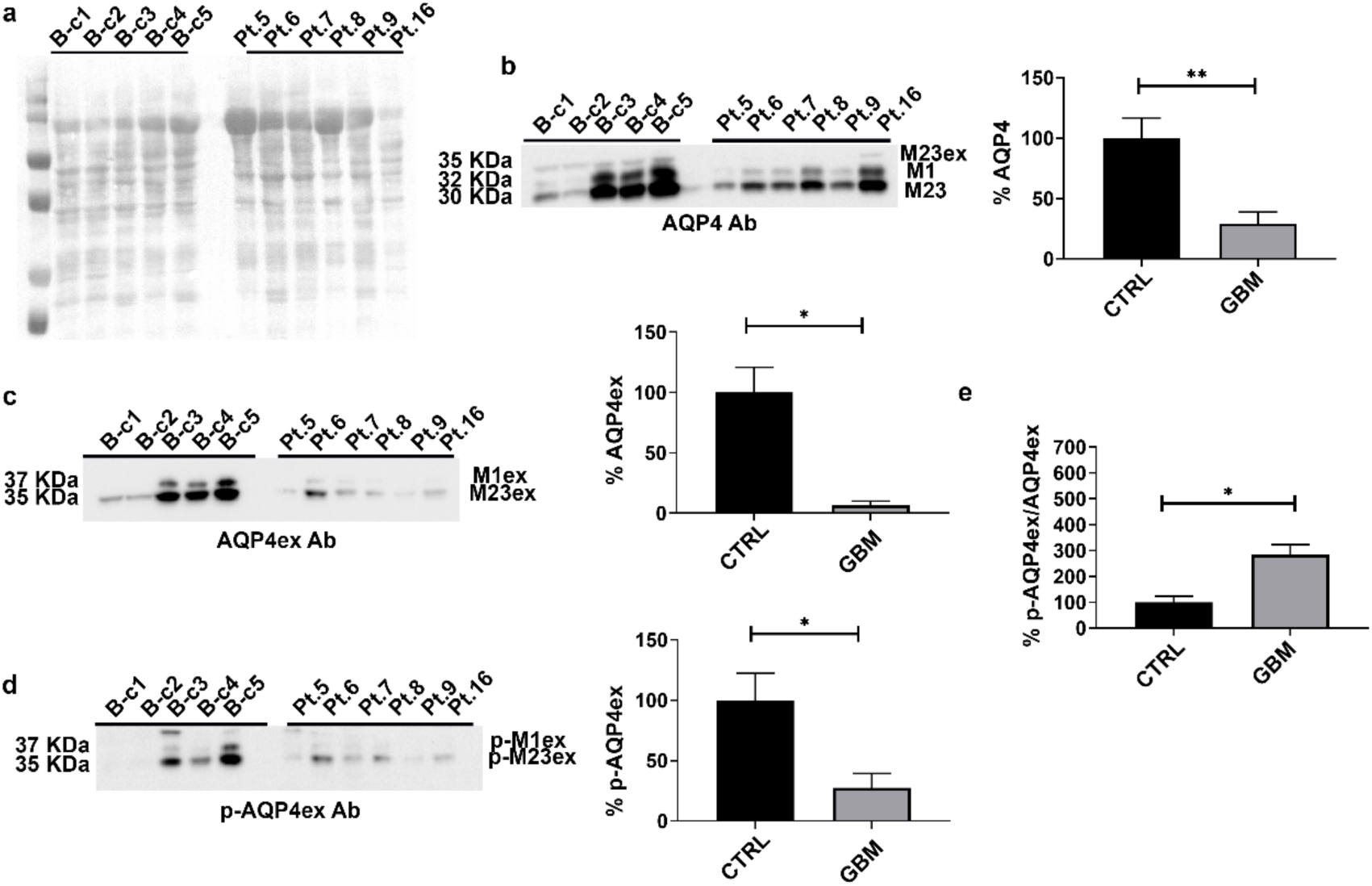
Quantitative analysis of AQP4 isoforms in GBM tumors. (a) Rouge Ponceaux staining of total proteins after transfer onto a PVDF membrane. (b-c-d) Immunodetection (left) and densitometric analysis (right) of AQP4 isoforms using antibodies for global AQP4 (b), AQP4ex (b), and p-AQP4ex (d), respectively. (e) Ratio between p-AQP4ex and AQP4ex and it shows an increase of AQP4ex phosphorylation levels in GBM compared to N (CTRL). (Asterisk indicated the student’s t-test significant differences for the comparison with basal level, *p < 0.05, **p<0.001). Brain CTRL (B-c1,2,3,4,5 n=5), GBM (n=6)

### AQP4 involvement in PA peritumoral edema

PA patients were divided into two groups according to the presence (EI >1) or not (EI=1) of peritumoral edema. Figure 5 (top) shows MRI scans representatives of the two categories. We evaluated the possible correlation between AQP4 expression and the presence of edema in PA. Immunoblot analysis (Fig. 5 bottom) revealed a moderate, but not significant increase of AQP4 expression in EI=1 group compared to controls and to the EI>1 group. Surprisingly, pAQP4ex expression was strongly increased in EI=1 compared to the other category (EI >1). Thus, although the expression levels of AQP4 did not appear to be related to edema index, the phosphorylation status of AQP4ex was strongly upregulated in EI=1 category.

**Fig 5.**
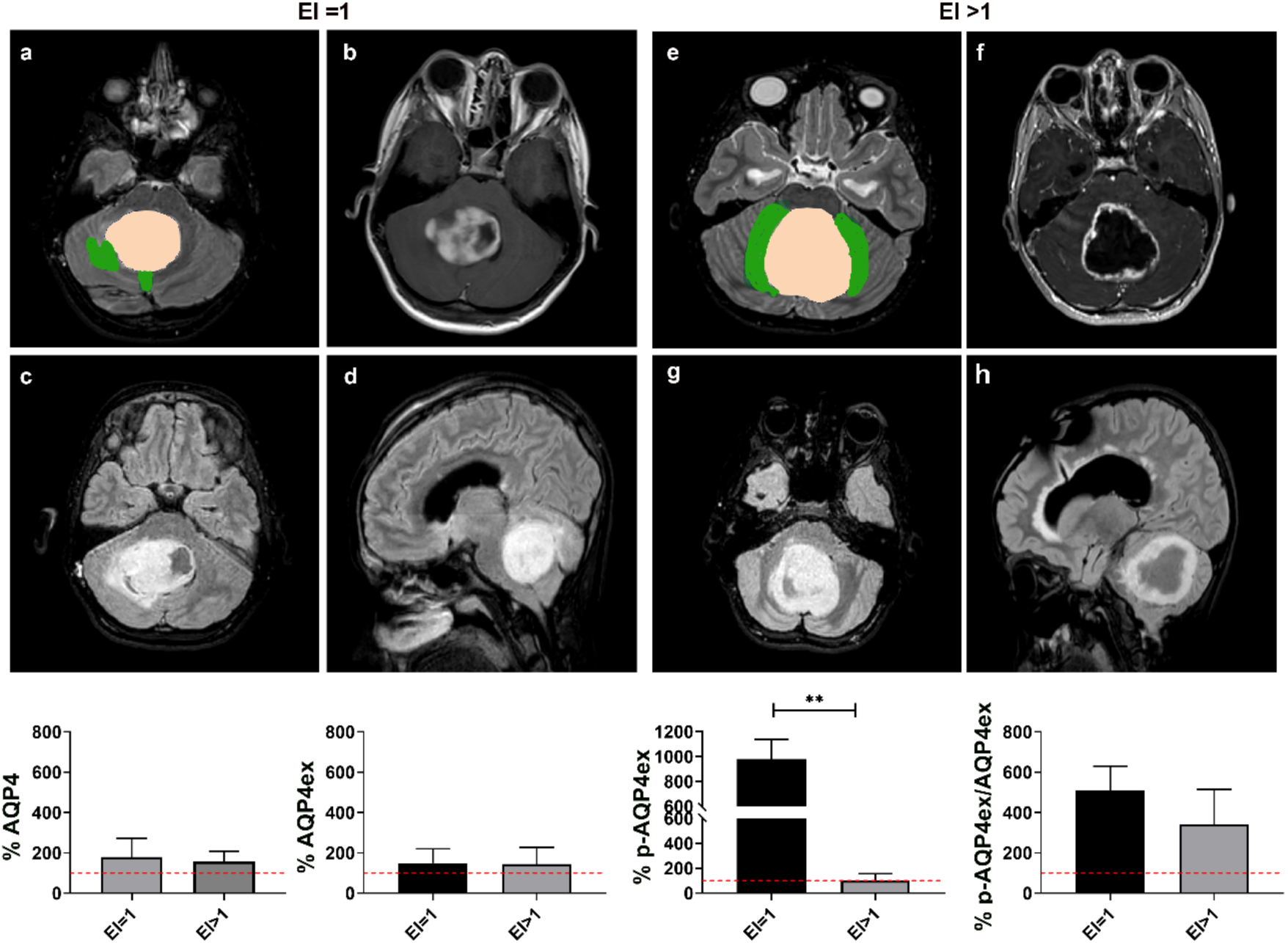
Evaluation of peritumoral edema and AQP4 levels in PA samples. (Top), MRI of PA samples biopsies of patient (Pt.8) in EI=1 group (left) and of a patient (Pt.7) in EI>1 group (right) in T2(a-e), T1 gadolinium-enhancement (b-f), FLAIR sequences (c-g); Tumor area was segmented in pink and edema was segmented in green in the T2 MR scan (a) for EI=1 and in the T2 MR scan (e) for EI >1. (bottom) AQP4 isoform expression level in the two EI categories. Dotted red line represents the controls set at 100%. **p < 0.001: Student t-Test; (N=5 for each EI category)

### Blood Brain Barrier is differently affected in PA and GBM tumors

To evaluate the BBB damage associated with glioma development we took advantage of the intraoperative use of sodium fluorescein (SF) during resection of both GBM [1, 2, 23] and PA [9].

We measured SF levels in PA T samples and compared them to N samples from GBM patients (Fig. 6a). Surprisingly, the SF concentration was found significantly higher in PA T samples to controls. Probably, this high level was related to the hypervascularization and peculiar characteristics of blood vessels in PA (see Figure 1) [27]. As expected, SF levels in GBM T samples were very high compared to unaffected tissues as previously reported by us [37] and other authors [35].

To evaluate the degree of endothelial proliferation we performed IF of VEGF a marker of hypervascularity, frequently associated with peritumoral BBB breakdown. Interestingly, VEGF was highly expressed in GBM T samples, while it was not detected in PA T biopsies, neither in controls (Fig.6b), indicating no gross abnormalities of BBB in PA and controls compared to GBM, as previously reported [12].This is consistent with the conclusion that the increased SF in PA is not related to BBB alteration but to the number of blood vessels and that GBM are fast malignant brain tumors that develop new blood vessels and express VEGF [24, 45].

**Fig. 6.**
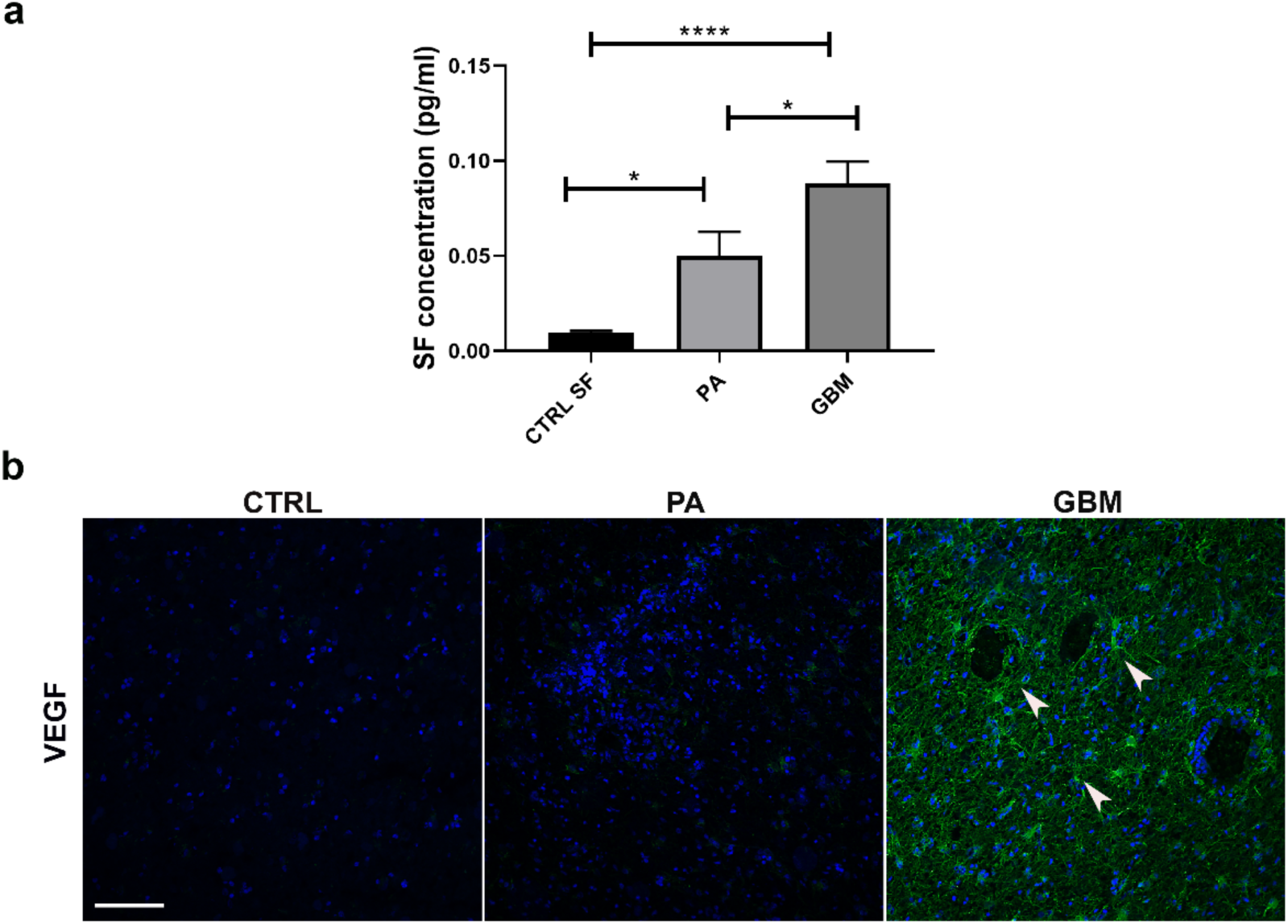
BBB evaluation in PA and GBM. (a) SF concentration is higher in PA T samples (n=5) compared to controls (CTRL, n=7), and highest in GBM T samples (n=9). One-Way ANOVA; post-test Tukey; *p<0.05; ****p<0.0001). (b) Immunofluorescence detection of VEGF in PA and GMB T samples. Arrowheads indicate strong VEGF expression in likely reactive astrocytes. Cell nuclei were stained with DAPI (in blue). Scale bar 300 µm

### Glial Fibrillary Acid Protein (GFAP) expression is increased in Glioma

To investigate whether and to what extent GFAP protein expression was altered in PA and GBM, IF and immunoblot experiments were performed. GFAP immunofluorescence signal was strongly increased in both PA and GBM T samples compared to controls (Fig. 7a) as a consequence of the strong gliosis, especially around blood vessels, occurring during glioma development.

To evaluate this last point, immunoblot detection of GFAP isoforms (α, δ, κ) was performed on both tumor categories (Fig. 7b, 7c). All GFAP isoforms were found up-regulated in both GBM and PA T samples compared to controls, confirming the strong reactive gliosis associated with tumor growth [8, 39]. Furthermore, the ratios between GFAP isoforms particularly (GFAPk/GFAPα) and (GFAPδ/GFAPα) were specifically analyzed since changes in these isoform ratios are implicated in the regulation of genes involved in tumor aggressiveness and malignancy [36]. Both GFAPk/GFAPα and GFAPδ/GFAPα were elevated in PA T samples compared to controls. Surprisingly, in GBM T samples the isoform ratios did not change substantially, likely due to a low number of samples (n=7) and the significant heterogeneity in GFAP expression observed in the analyzed GBM. Overall, the expression of GFAP was approximately three times higher in T samples of PA compared to GBM (Fig.7d).

**Fig. 7.**
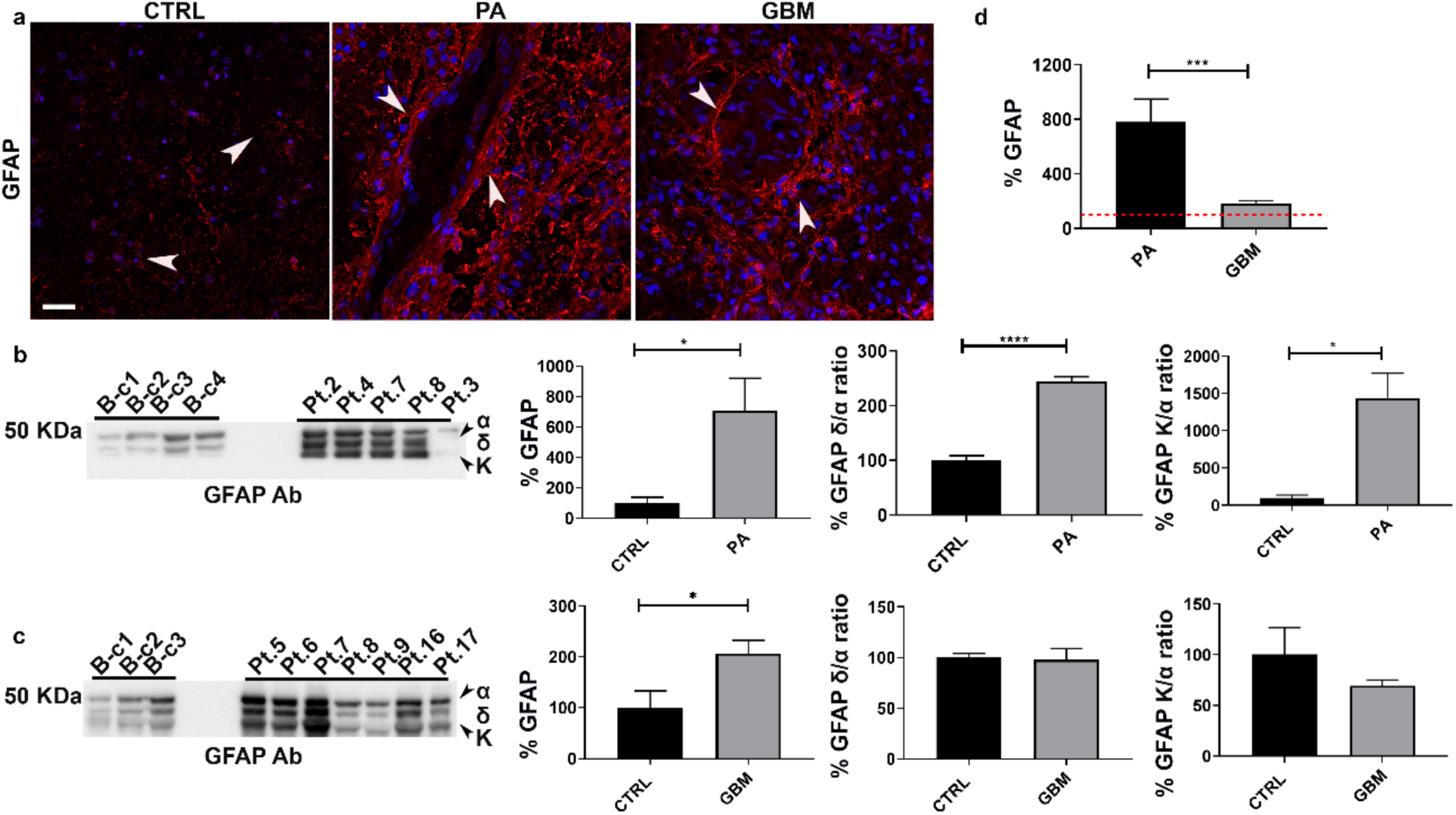
GFAP expression in PA and GBM. (a) Immunofluorescence in PA T samples (Pt.1) compared to control (CTRL) and GBM (Pt .5) T samples. GFAP staining (red) is different and higher in PA and GBM T samples compared to control (as indicated by white arrowhead). Cell nuclei were stained with DAPI (in blue). Scale bar 300 μm. (b-c) Immunodetection (left) and relative densitometric analysis (right) of global GFAP isoforms levels (α,δ,k indicated by black arrowheads) and rhe ratio between (GFAPk/GFAPα) and (GFAPδ/GFAPα) in PA (b) and GBM (c). (d) GFAP expression in PA (n=5) and GBM (n=7) T samples compared to control (CTRL, n=4) (represented as dotted red line seats at 100%) (*p < 0.05, ***p<0.001, ***p<0.0001: Student t-Test)

## DISCUSSION

### The Role of AQP4 Isoforms in PA and GBM

Our study offers critical insights into the role of AQP4 isoforms, particularly the extended isoform AQP4ex, in the context of gliomas, specifically PA and GBM, two types of brain tumors with markedly different biological behaviors and clinical outcomes. These findings are significant for understanding the mechanisms underlying edema formation and astrocytic responses, which vary considerably between these two tumor types.

In PA, we observed a significant increase in AQP4 expression, including AQP4ex. Despite this upregulation, AQP4 was found to be mis-localized, deviating from its typical perivascular localization at the astrocytic end-feet [10]. This pronounced mis-localization of AQP4 in PA mirrors findings in AQP4ex-KO mice, where the absence of AQP4ex results in the widespread mis-localization of the canonical AQP4 isoforms [21]. In the AQP4ex-KO mice, this mis-localization is associated with lower basal brain water content and impaired waste clearance, notably reduced amyloid-β (Aβ) drainage (*Abbrescia et al,.2024 submitted*). The parallels between AQP4 mis-localization in PA and AQP4ex KO mice suggest that proper AQP4 localization is crucial for maintaining water homeostasis and facilitating efficient waste clearance. This has significant implications for understanding the pathophysiology of PA, where despite the increase in total AQP4 levels and AQP4ex, the mis-localization of AQP4 could lead to functional impairments similar to those observed in the KO model. However, the increased amount of AQP4 in PA appears to partially compensate for the effects of mis-localization, as evidenced by the lack of significant edema typically associated with this tumor type. This suggests that the upregulation of AQP4 in PA, while not ideally localized, may still contribute to maintaining fluid balance, albeit less efficiently than if it were properly polarized. The minimal edema observed in PA, despite AQP4 mis-localization, could also be influenced by other factors such as a relatively intact blood-brain barrier (BBB) and a tumor microenvironment that does not promote extensive fluid accumulation.

AQP4 is capable of forming highly ordered two-dimensional structures in the cell plasma membrane called OAPs (Orthogonal Arrays of Particles)[40], the size of which depends on the ratio between the two canonical isoforms M1 and M23, as well as the presence of AQP4ex [6, 18, 21]. A recent study reports that overexpressing AQP4ex in mice reduces the formation of OAP [17] without affecting AQP4 polarization, suggesting that OAP formation is finely regulated by AQP4ex levels. Since in PA the overexpression of AQP4ex does not prevent AQP4 miss-localization, one possible explanation is that proteins involved in the AQP4 anchoring (i.e. syntrophin, dystrophin, etc) are also affected in PA. Another possibility is that level of AQP4ex phosphorylation in PA may play a role in the stability of AQP4 at the perivascular pole. Future research, including the generation of new transgenic animal mouse models, will be essential in elucidating these aspects.

### Impact of AQP4 Mis-localization on Water Homeostasis and Waste Clearance

In AQP4ex-KO mice, AQP4 mis-localization is associated with increased basal brain water content and significantly reduced efficiency in waste clearance. This finding is particularly relevant when considering its implications for PA. Although PA exhibit increased AQP4 expression, the mis-localization of this protein could similarly affect water dynamics and waste removal, though the extent of these effects may differ due to the compensatory increase in AQP4 levels.

The impaired waste clearance observed in AQP4ex-KO mice, particularly the reduced Aβ drainage, highlights the importance of proper AQP4 localization in facilitating the clearance of interstitial solutes. In PA, while there is no direct evidence of reduced waste clearance akin to what is observed in the KO model, the mis-localization of AQP4 could theoretically impair similar processes, potentially contributing to the long-term development of neurodegenerative conditions if not adequately managed.

The data suggest that AQP4ex plays a crucial role in anchoring AQP4 to the astrocytic end-feet, essential for maintaining brain water homeostasis and ensuring efficient solute clearance. In the absence of AQP4ex, as seen in the KO model, or in cases of mis-localization, as observed in PA, these critical functions are affected, leading to altered brain water content and impaired waste clearance mechanisms.

A novel aspect is the role of AQP4ex phosphorylation in those different tumors. While in GBM low AQP4ex production and low phosphorylation could trigger a reduction in global AQP4 expression, in PA pAQP4ex levels dramatically increases. This suggests that phosphorylation may regulate the expression and the stability of AQP4 at the BBB level, enhancing its role in modulating edema formation in a manner specific to tumor type.

The increase in the canonical AQP4 isoforms in PA may be related to the escape of the readthrough mechanism and/or vascular hyperproliferation which could activate astrocytes around blood vessels. Indeed, PA show reactive astrocytes especially in the acidophilic structures enriched with Rosenthal fibers around blood vessels [11, 46].

### GFAP Expression and Reactive Astrogliosis

Our findings on GFAP expression provide important insights into astrocytic responses in PA and GBM. GFAP, a well-known marker for astrocytes, is typically upregulated during reactive astrogliosis, a common response to brain injury or disease. While some studies have reported that GFAP expression correlates with tumor grade [5], others have observed an opposite trend [15]. In our study, we observed a significant increase in GFAP levels, independent of glioma grade, suggesting a strong reactive gliosis associated with tumor growth. The elevated GFAP expression in PA may be linked to vascular hyperproliferation, which could activate astrocytes around blood vessels. Notably, reactive astrocytes were observed around blood vessels, particularly in Rosenthal fibers, which are characteristic of PA. [11, 46]. These fibers are thought to contribute to the activation of astrocytes and the consequent upregulation of GFAP in PA [11, 14, 32].

GFAP expression seems to be influenced by the degree of cellular differentiation: lower grade astrocytomas tend to have higher GFAP levels compared to more undifferentiated tumors like GBM [31] characterized by undifferentiated regions and prone to dedifferentiation [26].

In our study, we found a significant increase in GFAP expression in both PA and GBM, with much higher levels in PA. This upregulation includes specific increases in the GFAPκ and GFAPδ isoforms, particularly in PA. These findings are consistent with previous studies, underscoring the role of GFAP isoform expression in differentiating astrocytic tumors [33, 38].

In PA, the significant increase in GFAP expression is closely linked to the presence of Rosenthal fibers—elongated, eosinophilic structures found within astrocytes. These fibers, characteristic of PA, suggest long-standing reactive astrocytic processes. The high GFAP levels likely reflect a pronounced state of reactive gliosis, where astrocytes actively respond to the tumor microenvironment, potentially contributing to the relatively benign nature of these tumors by stabilizing the surrounding tissue and limiting tumor spread. Conversely, GBM also show increased GFAP expression, though without the association of Rosenthal fibers. In GBM this elevation likely represents a generalized reactive gliosis driven by the aggressive and invasive nature of the tumor, rather than a protective or stabilizing reaction. The absence of Rosenthal fibers in GBM suggests that the increase in GFAP in these tumors does not confer the same effect seen in PA, but rather reflects an astrocytic response to the rapidly expanding and infiltrative tumor mass.

## Conclusion

In conclusion, our study highlights the dual and opposing effect of AQP4 and its perivascular isoform AQP4ex in PA and GBM, reflecting the complex and tumor-specific nature of astrocytic responses in gliomas. The distinct behaviors of AQP4 and GFAP in PA and GBM underscores the need for tumor-specific therapeutic approaches. In PA, the upregulation and mis-localization of AQP4, along with GFAP-associated Rosenthal fibers, suggest that modulating these responses could benefit the management of low-grade gliomas, limiting their growth and infiltrative capacity. Conversely, in GBM, therapeutic efforts might focus on restoring AQP4ex expression and its proper polarization while modulating GFAP to reduce reactive gliosis. AQP4 holds potential as a biomarker for glioma progression, offering insights into water homeostasis and astrocytic function that could inform future therapeutic strategies aimed at improving patient outcomes in gliomas.

## DECLARATIONS

### Author Contributions

OV performed quantified immunoblots and immunofluorescence experiments. RM, LDG, MTB and FS performed the biopsies. GI, and AD performed histopathological tissue analysis. DSZ performed MRI. PA, CP and GS performed tissue preparation, freezing and analysis. GPN, AF and FS designed the study. OV and AF wrote the manuscript.

All authors read and approved the final manuscript.

### Availability of data and materials

All data generated or analyzed during this study are included in this published article

### Ethics approval

This research was performed in compliance with institutional guidelines and approved by the appropriate institutional committees (project. n.6898). All applicable international, national, and/or institutional guidelines for the care and use of animals were followed.

### Consent for publication

Not applicable.

